# Impaired baroreceptor reflex sensitivity in pelvic suspension rats is associated with sympathovagal imbalance but not with carotid artery structural change

**DOI:** 10.1101/2022.09.02.506352

**Authors:** A Bhatnaagar, KK Deepak, A Roy, AK Anil, B Sharma, A Singh, RK Netam

## Abstract

**Introduction:** In space, cephalad shift of body fluid equalizes the blood pressure throughout the body leading to various cardiovascular disturbances in space and upon return on earth. Effects of microgravity on humans can be simulated on earth by hindlimb unloading (HU) rat models l. This study was planned to examine the Effect of 15 days hindlimb unloading on baroreflex sensitivity, Herat rate variability (HRV) and carotid artery structure in rats.

**Material and methods:** 18 male Wistar rats weighing 250-300 gm were randomly divided into control and hindlimb unloading groups and housed in separate cages for 15 days. Body weight, food and water intake was measured every day. The pelvic suspension method was used to simulate effects of microgravity. To record HRV, ECG was recorded in lead II configuration on the 15th day. Then blood pressure was recorded directly from the carotid artery. Linear regression of arterial pressure and RR interval used to derive the Baroreflex sensitivity. Vessel wall thickness, intimal thickness, smooth muscle cell layer thickness &amp; luminal diameter were compared after H&E staining. Ultrathin sectioning and counter-staining was done for electron microscopy and Changes in endothelial thickness, elastic lamina, and deeper artery structures were observed. Statistical analysis was done using Graphpad Prism (version 9.0). Unpaired t-test, Mann Whitney and 2 way ANOVA were used according to groups and distribution of parameters.

**Results:** We found significant differences in Food intake (p □0.001), Body weight (p □0.001), Heart rate (p=0.031), BRS (p= 0.026), R-R interval (p=0.0246), heart rate variability parameters {SDSD (p=0.0001), RMSD (p= 0.0001), VLF(p=0.001),HF (p=0.0054), LF/HF Ratio (p= 0.0001)}. We did not found significant changes in Water intake, MAP, SBP, DBP, Vessel wall thickness, Luminal Diameter, Intimal thickness, Smooth muscle thickness, in H&E staining and in electron microscopy.

**Conclusion:** Chronic microgravity simulation using pelvic suspension showed a decrease in BRS after 15 days of HU; however 15 days of HU was not insufficient to produce any structural changes in the carotid artery. BRS changes were accompanied by change in vascular vasoconstriction and vasodilation properties which were due to alteration in the sympathetic and parasympathetic outflow.

## Introduction

Due to the earth gravity, the human body in an upright position on the earth’s surface has differential pressure distribution throughout the body. This evolutionary change of differential pressure distribution maintains the proper blood flow in different organs of the body. However, the exposure to microgravity during spaceflight and space stations vanishes this differential pressure distribution and causes major alterations in the cardiovascular system that results, fluid loss, fluid redistribution, decrease stroke volume, decreased aerobic capacity, and cardiac atrophy. (1,2) These alterations pose difficulties in space operations and adaptability after returning to Earth. Due to limitations of human subjects, laboratories like NASA prefer to study microgravity simulation on rat models. Results of these studies can then be used to develop strategies and technologies to protect astronauts’ cardiovascular health during prolonged space missions. (3-4)

Hind limb unloading (HU) models successfully replicate a few of the microgravity effects, including cephalic fluid shift, BP changes, bone loss, tachycardia, weight loss, and muscle atrophy. Acute and chronic microgravity studies have been conducted with HU models at various unloading inclinations (20 to 45º). Acute microgravity simulation using tail suspension showed a decrease in BRS, which returned to baseline after 48 hours. (5) The chronic microgravity simulation using tail suspension, on the other hand, resulted in a sustained decrease in BRS. (6). BRS is mainly affected by the structural modification of carotid artery/aortic arch and autonomic responses to the change in blood pressure. Baroreflex is the main autonomic reflex which regulates blood pressure in normal range.(7) Afferent components of baroreflex arch includes baroreceptors present aortic arch and common carotid artery and afferent nerve (glossopharyngeal & vagus nerve) which takes signal to NTS and VMC in medulla oblongata.(8) Sudden increase in blood pressure led to stretching aortic arch and common carotid artery which stimulates baroreceptors, then through afferent nerve signal goes to brain.(9) Hemodynamic burden due to microgravity exposure can lead to Structural modification of major arteries such as change in proliferation of smooth muscle & cellular and extracellular matrix. (10) The efferent component of Autonomic reflex arc is sympathetic and parasympathetic input to heart and blood vessels which controls the heart rate and blood pressure in response to change in BP.

Thus, BRS is primarily influenced by the mechanical characteristics of the major arteries and the tone of the sympathetic and parasympathetic nervous systems; no prior research has compared BRS with the structural alterations of the carotid artery and changes in autonomic tone in the hind limb suspension rat model Thus, the aim of this work is to compare the BRS, autonomic tone, and structural alterations of the carotid artery following 15 days of HU with control rats.

## Materials and methods

This research was conducted at the AIIMS, New Delhi, in the Physiology Department. The AIIMS Institutional Animal Ethics Committee granted us ethical permission (File No. 153/IAEC-1/2019). The National Institute of Health’s (NIH) Guidelines for the Care and Use of Laboratory Animals (NIH Publication no. 85723, amended 1996) were followed in conducting the research. Eighteen mature male Wistar rats weighing 250 grams were obtained from the AIIMS, New Delhi’s Central Animal Facility. Ad libitum access to clean drinking water and laboratory food pellets were supplied to every rat kept in its own polypropylene cage. Each animal was kept in a room with an ambient temperature of 24±2°C, a relative humidity of 50–55%, and a light–dark cycle of 12 hours. After the initial acclimatization period rats were randomly divided into 2 groups; control and hind limb suspension.

### Hip Suspension of rats

Hind limb suspension rats were suspended in the cage using a plastic wire which was passed through the brace. Suspension cage was fabricated similar to the cage used by Morey-Holton with certain modifications. (11) The metallic brace covered with leucoplast were curved in the shape of the pelvic brace. Once the pelvic brace was ready, the rat was anesthetized using ketamine & xylazine and it was fitted into the brace. The brace was tightened to ensure that the rat was not able to escape. (Figure1) Rats were allowed to acclimatize to the brace for 7 days, and then rats were suspended at an angle of 30 degrees by passing plastic insulated wires through the brace for 15 days. (12)

**Figure 1:**
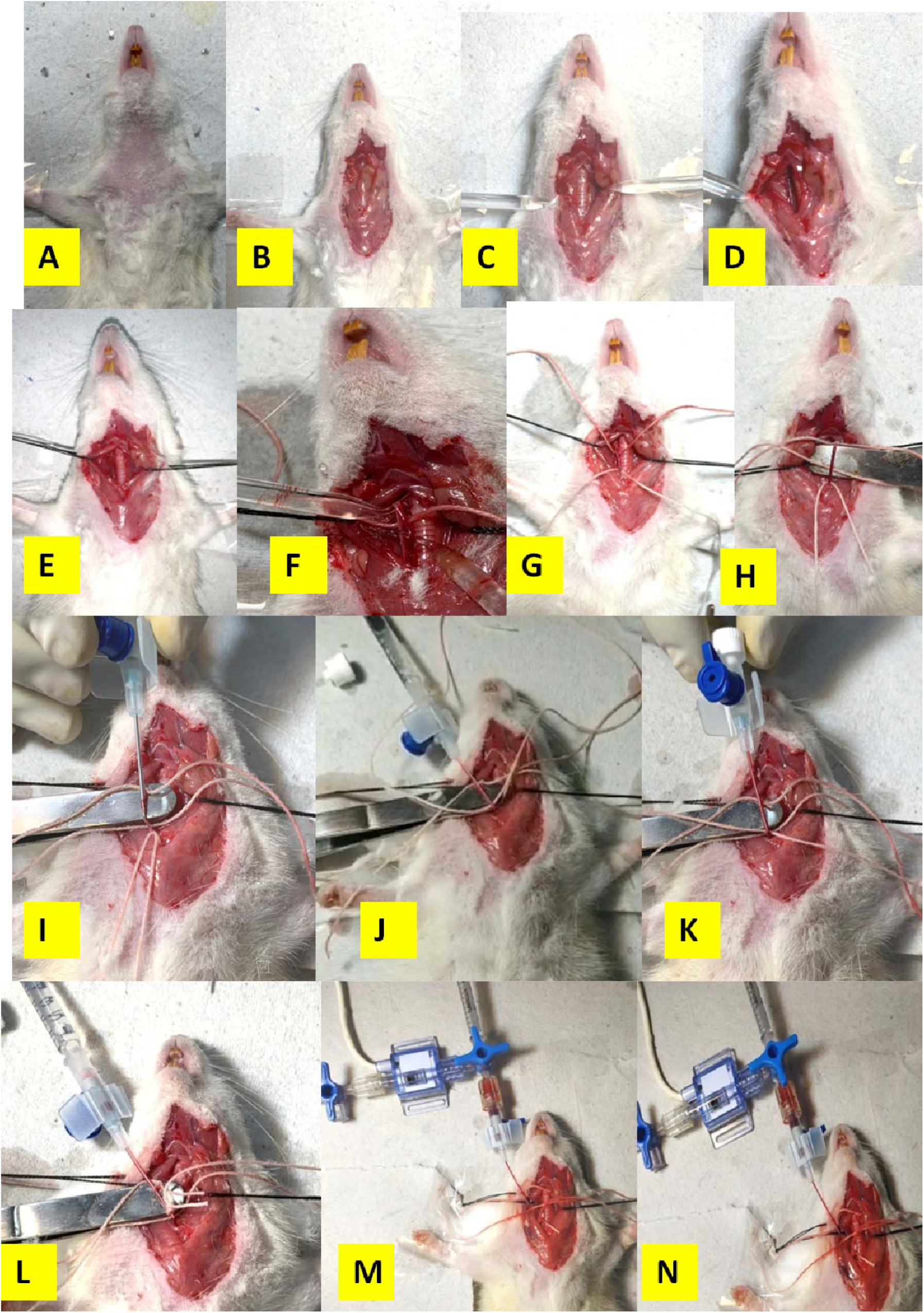
Steps of invasive blood pressure recording from carotid artery in rat. **Step A:** Shaving of chest of the rat. **Step B:** Thymus and other glands visualization after removal of skin in the chest. **Step C:** Visualization of trachea. **Step D:** Visualization of vagus nerve and carotid artery in the side of trachea. **Step E:** side muscles are sutured to expose the surgical area. **Step F:** Vagus nerves were dissected out from carotid sheath. **Step G:** 2 Threads were passed behind the carotid artery. **Step H:** A support is given from behind to the carotid artery for cannulation. **Step I:** A nick is made by 16 gauze venflon. **Step J:** Trocar of venflon is removed and heparinised saline was injected in artery to prevent clotting. **Step K:** Venflon is closed by cap and cannulation is secured by a knot. **Step L:** heparinised saline was injected in artery to prevent clotting. **Step M:** fluid filled pressure transducer was attached to cannula and baseline BP was recorded. **Step N:** different dose of phenylepinephine and nitroprusside was injected and change in mean arterial pressure, R-R interval & HR were recorded for the same.

### Recording of Body weight, food intake, and water intake

Rats’ body weight was measured daily using a weighing machine (sensitivity: 10 gm, range: 0-500 g). Standard rat chow was supplied to the rats, and weighing techniques were used to record their food intake. 100 ml of water was provided to every rat per day at 10 am, and the amount of water left over was noted the next day at 10 am in order to determine the water consumption.

### Baroreflex sensitivity recording

After 15 days of suspension, the rats were given intraperitoneally urethane (1.5 g/kg) to induce anaesthesia, and the depth of the anaesthesia was assessed by the rats’ withdrawal reflex. After shaving the hair from the anterior region of the neck, the rat was then put ventrally on the surgical table, and the surgical site was cleansed with betadine. (Figure 1) To record heart rate, ECG leads were attached in lead II configuration. 0.5 cm a midline skin incision was made by a surgical scissor, followed by a blunt dissection of glands, subcutaneous tissue, and omohyoid muscle was done by two glass seekers. The carotid artery was identified as a big, palpating artery with a vagus nerve on the side of the trachea. **(Figure 1)** The vagus nerve was carefully isolated from the artery, and the carotid artery was cannulated with pre-heparinised 24-gauge venflon. To prevent clotting, 0.5 ml of heparinised saline was slowly flushed into the heart immediately after cannulation. The cannula was then attracted to the fluid-filled pressure transducer (Power lab, AD Instruments). Blood pressure and heart rate were measured at rest. After 30 minutes of baseline recording, blood pressure and heart rate responses were randomized to four doses of epinephrine (10, 20, 40 & 60 μg /kg) and sodium nitroprusside (5, 15, 20 & 35 μg/kg). Following each dose, the blood pressure and heart rate were given enough time to recover to baseline. (13)

### Hear rate variability recording

Before recording BRS, HRV recording was done, for that chest of the rats was properly shaved and cleaned then rats were made to wear custom build chest jacket as described by Pereira-junior et al 2010 which already had 3 prefixed platinum electrodes. Before starting the procedure conductive gel was applied for the proper contact between skin and electrode. After rat was made to wear jacket, rats were restrained in the retainer, and electrodes were connected to the power lab (ADInstruments, Australia) by wires. Rats were made to acclimatize in restrainers for 30 minutes and lead II EEG was recorded. (14)

### Histology

After blood pressure recording, rats were immediately scarified and perfused with normal saline. Then the carotid artery was dissected out and cannulated and pressurized with oxygenated Kreb solution at 37□C in 100 mmHg and fixed in a paraformaldehyde solution for fixation. Fixed Samples were stored in a 4°C for 24 hours. Stored tissues were then sequentially transferred to sucrose solutions in the next 2 days, followed by 8 micrometer thick sectioning by cryostat microtome (Leica, Germany) and staining by hematoxylin and eosin. (**Figure 2**) The imaging was done using the NIS Element basic research software. Vessel wall thickness, intimal thickness, smooth muscle cell layer thickness &amp; luminal diameter were assessed by the Fiji ImageJ software.

**Figure 2:**
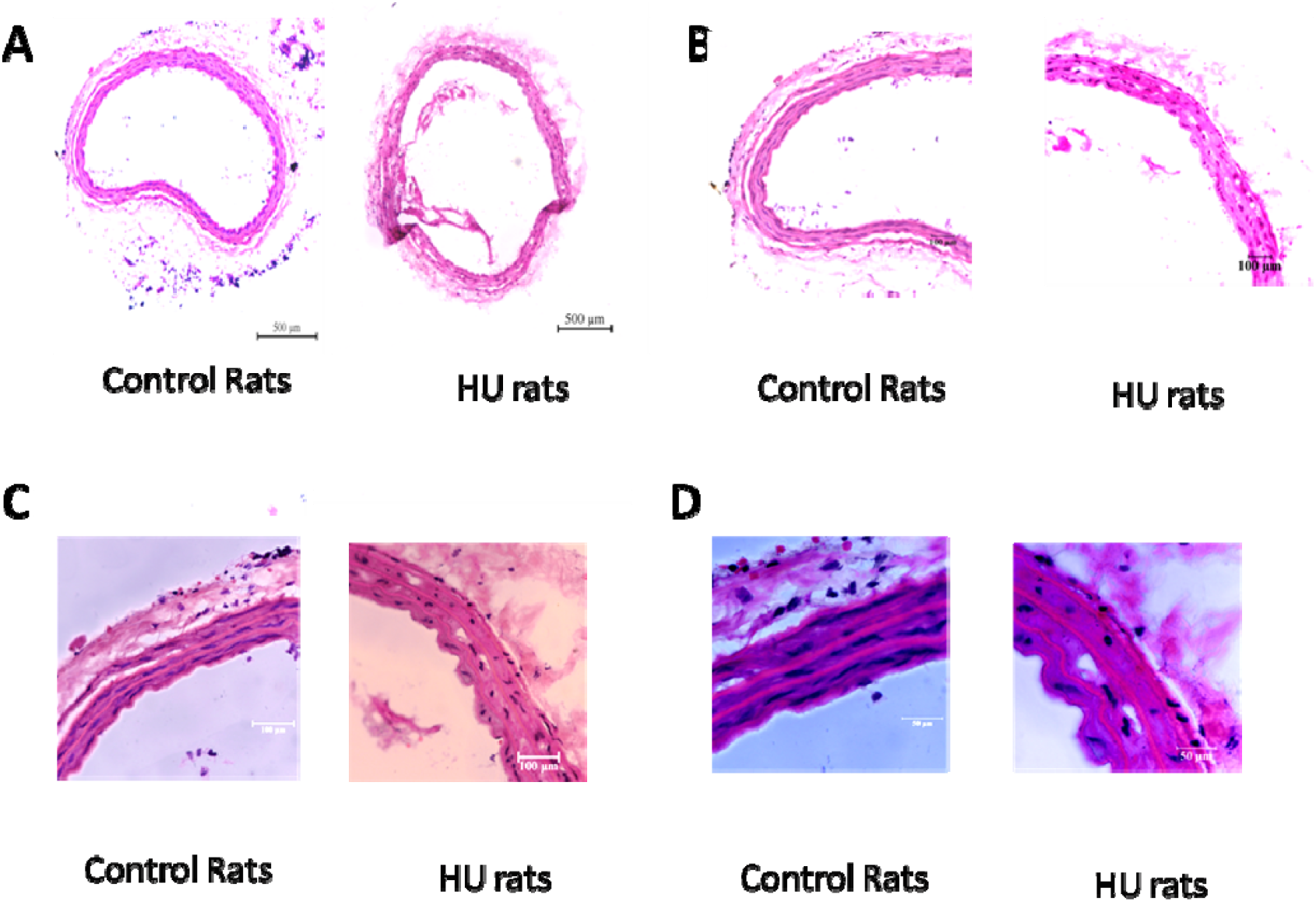
Carotid artery histology in different magnification (H & E staining) Figure 2: comparison of carotid artery structure (H & E staining) after 15 days of pelvic suspension. Representative histological sections of the pelvic suspended and control group at different magnification Panel A:4x, Panel B:10x, Panel C:40x, Panel D:100x.. The imaging was done using the NIS Element basic research software and Fiji ImageJ software were used to calculate Vessel wall thickness, Luminal Diameter, Intimal thickness and smooth muscle layer thickness the .N=6-9 Control and 6-9 pelvic suspension rats.

### Electron microscopy

The carotid artery was dissected and placed in the mixture of 2% Paraformaldehyde and 2.5% glutaraldehyde in 0.1 M phosphate buffer saline at 4°C for 24 h. Secondary fixation of the artery was done in the mixture 1% osmium tetroxide and 1.5% potassium ferrocyanide at 0°C for one hour. After that tissue was transferred for an additional one hour in 1% OsO4 in 0.1 M sodium cacodylate. Then the tissue was dehydrated with different concentrations of ethanol. Tissue infiltration was done by propylene oxide: Durcupan (1:1), then tissue embedding by Durcupan for 48h. To select areas for ultrathin sectioning (70-80nm), a semi-thick tissue section (500nm) was first cut and sections were microscopically evaluated. Tissue sections with good visibility of all the vessel characteristics were then subjected to ultrathin sectioning. Ultrathin sections (100nm) were cut by Ultracut S microtome (Leica) and counter-staining was done by lead citrate, then sections were observed under a Tecnai G2 20 electron microscope (Fei Company, The Netherlands). Images were digitally acquired at 750-2250 X magnification by a charge-coupled device (CCD) camera. Representative image of electron micrograph is shown in **Figure 3**.

**Figure 3:**
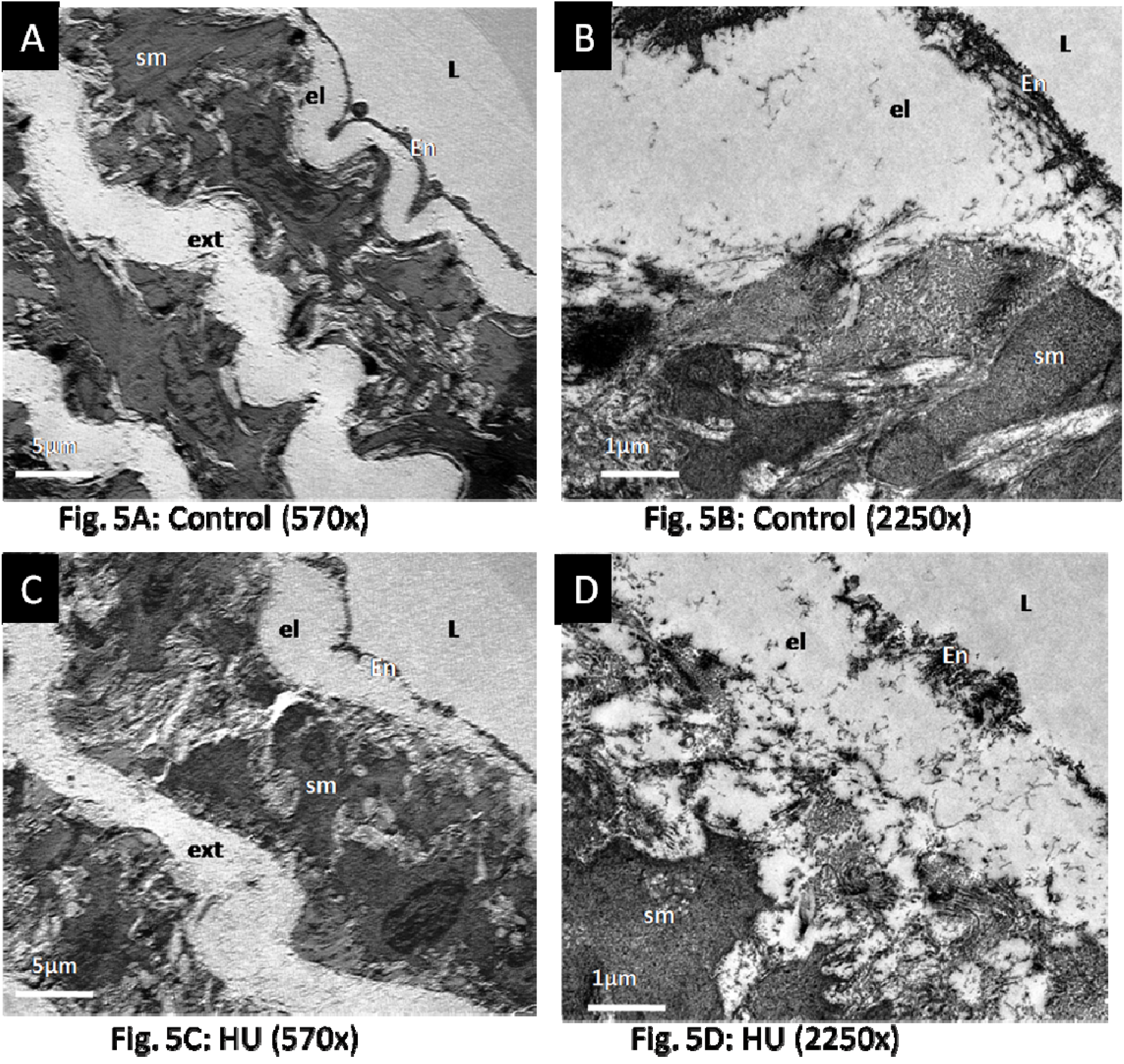
comparison of Electron microscopy micrograph of pelvis suspended rats with control rats. Panel A: Electron microscopy micrograph of a normal rat showing normal endothelial layer (EN), internal elastic lamina (IEL), smooth muscle layer (SM), external elastic lamina (EEL) at 570X and Panel B: at 2250X Panel C: Electron microscopy micrograph of a pelvic suspension Rat’s carotid artery at 570X and panel D: 2250X Unpaired tests test showed no significant change in EN, IEL, SM and EEL. P< 0.05 *; <0.01 **, <0.001 ***, <0.0001 **** control=6-9 and pelvic suspension=6-9

### Statistical analysis

Statistical analysis was done using Graphpad prism (version 9.0). Data was first tested for distribution using the Shapiro-Wilk normality test. According to distribution unpaired t test or Mann Whitney were used. To calculate BRS, different doses of phenylepinephine (5,10, 15 &amp; 20 µgm/kg) and sodium nitroprusside (6,12, 24 &amp; 48 µgm/kg) were given through carotid artery and HR rate and the MAP was recorded. The MAP and R-R Interval are plotted and the slope of the graph is calculated and taken as the Baroreflex sensitivity. To derive baroreflex sensitivity linear regression was done between RR interval and mean arterial pressure change. Parametric data is expressed as mean ± SD and non parametric data is expressed as median (Interquartile range). Body weight, Food intake, water intake and dose response curve for different doses of phenylepinephine and sodium nitroprusside were compared using 2 way ANOVA and post hoc comparison was done using the Bonferroni post test. P< 0.05 *; <0.01 **, <0.001 ***, <0.0001 **** control.

## Results

Changes in the bodyweight of both the groups of rats were compared at end of 1^st^ day, 5^th^ day, 10^th^ day, and 15^th^ day. There was a significant weight difference in between pelvic suspension group and the control group at 10^th^ day (p value □0.001) and 15^th^ day (p value □0.001). [(Baseline, C: 264.16±4.55 vs HU: 265.83±5.23)], [(Day 1, C: 263.33±4.95 vs HU: 261.67±4.77)], [Day 5, (C: 268.33±4.77 vs HU: 251.67±4.01)], [Day 10, C: 275±4.28: vs HU: 241.67±4.01) and [Day 15, C: 278.33±5.43: vs HU: 241.67±3.07). **(Figure 4A)**

**Figure 4:**
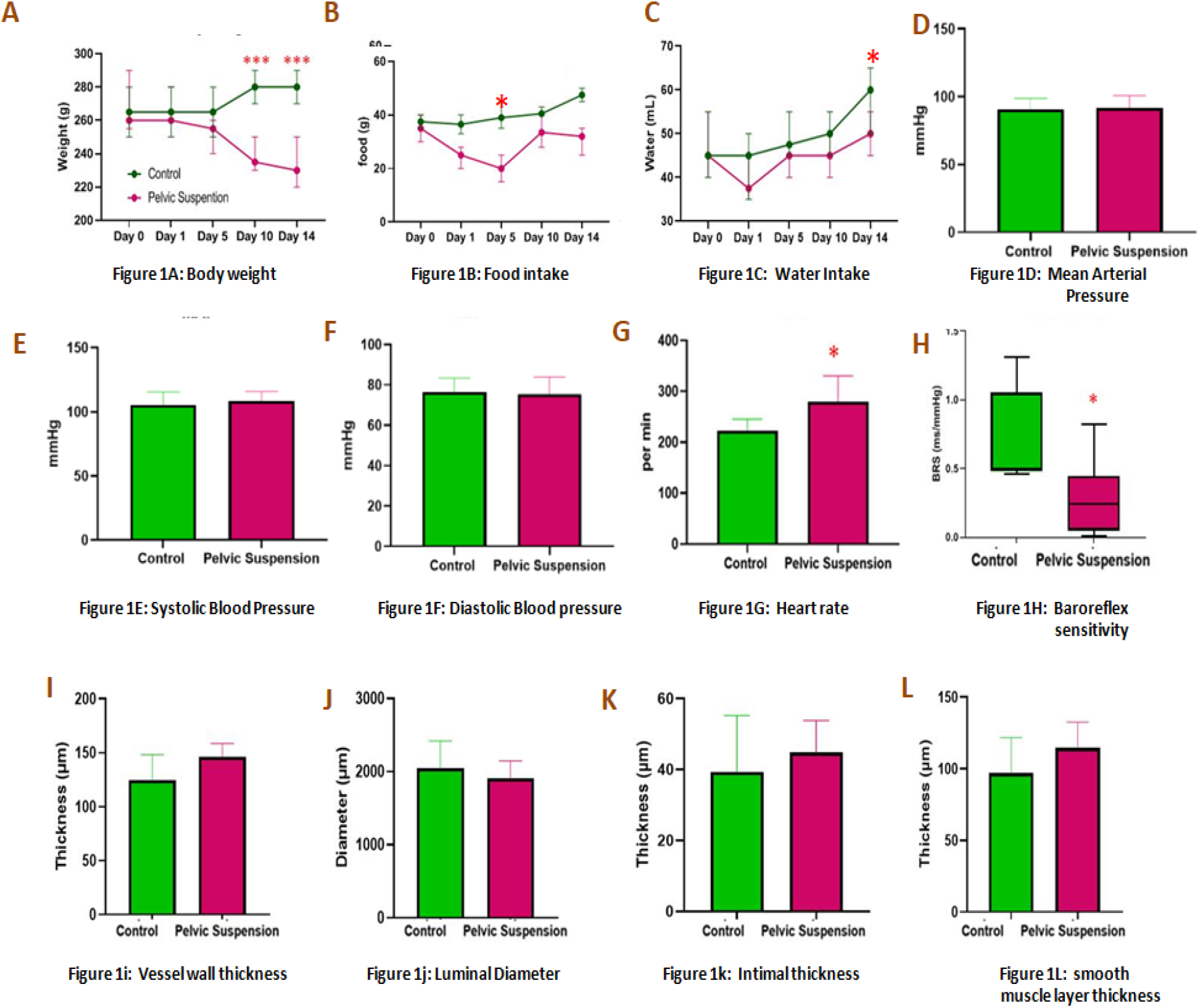
Graphical representation of *comparison of* between control and HU rats. **Figure4: comparison of body weight, food intake, water intake,, Mean arterial pressure, systolic pressure, diastolic pressure, baroreflex sensitivity, Vessel wall thickness, Luminal Diameter, Intimal thickness and smooth muscle layer thickness of carotid artery between control and pelvic suspension rats after 15 days.** **Figure4 A:** comparison of Body weight (Grams). Two-way ANOVA showed significant effects of interaction (F (4, 40) = 17.47 ; P = <0.0001) and Pelvic suspension F (1, 10) = 10.99 ; P < 0.0001), but not for day of experiment (P = 0.0078). The Bonferroni post test showed significant differences between control and pelvic suspension on days 10 and 14. * * P= 0.0010, **P < 0.0019; control=6 and pelvic suspension=6 **Figure4 B :** comparison of Food intake as gram/day. Two-way ANOVA showed significant effects of Pelvic suspension F (1, 10) = 21.37; P < 0.0009), but not for day of experiment (P = 0.0884). The Bonferroni post test showed significant differences between control and pelvic suspension on days5 *P < 0.0160 ; control=6-9 and pelvic suspension=6-9 **Figure 4 C :** comparison of water intake (mL). Two-way ANOVA showed significant effects of Pelvic suspension F (1, 10) = 11.75 ; P =0.0065) and time F (2.347, 23.47) = 13.89 P<0.0001. The Bonferroni post test showed significant differences between control and pelvic suspension on days 15 *P < 0.0160 ; n=Control: 6-9 and pelvic suspension: 6-9 **Figure 4D:** comparison of *comparison of* mean arterial pressure of pelvic suspension rats with control rats after 15 days of HU. BRS at 15th day. Unpaired T test; 2 tail with 95% confidence interval [C: 90.48±8.37mmHg vs Ex: 91.77±8.97mmHg (p=p=0.801)] ; n=Control: 6-9 and pelvic suspension: 6-9 **Figure 4E:** comparison of Systolic blood pressure of pelvic suspension rats with control rats after 15 days of HU. SBP at 15th day, Unpaired T test; 2 tail with 95% confidence interval [C: 105.6±10.02 mmHg vs HU: 108.4 ±7.5 mmHg (p=p=0.592)] ; n=Control: 6-9 and pelvic suspension: 6-9 **Figure 4F:** comparison of Diastolic Blood pressure of pelvic suspension rats with control rats after 15 days of HU. DBP at 15th day, Unpaired T test; 2 tail with 95% confidence interval [C: 76.43±7.04 mmHg vs Ex: 75.55± 8.3 (p=0.848)]. ; n=Control: 6-9 and pelvic suspension: 6-9 **Figure 4G:** comparison of HR of pelvic suspension rats with control rats after 15 days of HU. Unpaired T test; 2 tail with 95% confidence interval [C: 223.2±22.3/min vs HU: 279.9±0.75/min (p=0.031)] ; control=6-9 and pelvic suspension=6-9 **Figure 4H:** comparison of BRS of pelvic suspension rats with control rats after 15 days of HU. Unpaired T test; 2 tail with 95% confidence interval [C: 0.4957(0.4801-1.311) vs Ex: 0.2429(0.0499-0.4462) (p=0.026)]. ; n=Control: 6-9 and pelvic suspension: 6-9 **Figure 4I:** comparison of vessel wall thickness of carotid artery of pelvic suspension rats with control rats after 15 days of HU. Unpaired T test; 2 tail with 95% confidence interval [C: 0.4957(0.4801-1.311) vs Ex: 0.2429(0.0499-0.4462) (p=0.026)]. ; n=Control: 6-9 and pelvic suspension: 6-9 **Figure 4J:** comparison of Luminal Diameter of carotid artery pelvic suspension rats with control rats after 15 days of HU. Unpaired T test; 2 tail with 95% confidence interval [C: 0.4957(0.4801-1.311) vs Ex: 0.2429(0.0499-0.4462) (p=0.475)]. n=Control: 6-9 and pelvic suspension: 6-9 **Figure 4K:** comparison of Intimal thickness of carotid artery pelvic suspension rats with control rats after 15 days of HU. Unpaired T test; 2 tail with 95% confidence interval [C: 0.4957(0.4801-1.311) vs Ex: 0.2429(0.0499-0.4462) (p=p=0.47)]. n=Control: 6-9 and pelvic suspension: 6-9 **Figure 4L:** comparison of smooth muscle layer thickness of carotid artery of pelvic suspension rats with control rats after 15 days of HU. Unpaired T test; 2 tail with 95% confidence interval [C: 0.4957(0.4801-1.311) vs Ex: 0.2429(0.0499-0.4462) (p=0.189)]. n=Control: 6-9 and pelvic suspension: 6-9, P< <0.05 * ; <0.01 **, <0.001 *** ; <0.0001 **** 001 vs. Control

Difference in the food intake in both the groups of rats was compared at the end of 1^st^, 5^th^, 10^th^ and 15th day (Fig.2). There was significant less food intake (p value □0.001) in pelvic suspension group as compared to control group at 5^th^, 10^th^ and 15th day.). [(Baseline, C: 35±3.16vs HU: 30±4.49)], [(Day 1, C: 34±2.98 vs HU: 22.17±2.66)], [Day 5, (C: 36.17±8.61 vs HU: 19.17±3.76)], [10^th^, C: 37.33±8.57vs HU: 30±3.47] and [15^th^, C: 43.33±4.69 vs HU: 28.5±3.03] **(Figure 4B)**. Figure 4C shows a difference in the water intake in both the groups of rats at the end of 5th, 10th and 15th day. There was no significant difference in water intake in pelvic suspension group as compared to control group (p value □0.001).). [(Day 0, C: 43.33±5.05 vs HU: 42.5±8.22)], [(Day 1, C 40±4.75: vs HU: 35.83±3.59)], [Day 5, (C: 43.33±5.05 vs HU: 40±3.23)] and [10^th^ day {46.67±8.76 vs HU: 41.5±3.62) and [15^th^ day, (53.33±4.10vs HU: 50.83±2.44). **Figure 4C**

Difference in the MAP, SBP and DBP of both the groups of rats were compared at the end of 15th day respectively (Fig. 4D-F). There was no significant difference in between pelvic suspension group and control group at 15th day. MAP at 15th day [C: 90.48±8.37mmHg vs Ex: 91.77±8.97mmHg (p=0.801)], SBP 15^th^ day [C: 105.6±10.02 mmHg vs HU: 108.4 ±7.5 mmHg (p=0.592)], DBP [C: 76.43±7.04 mmHg vs Ex: 75.55± 8.3 (p=0.848)]. **Figure 4D-F** shows HR of both the groups of rats at the end of 15th day. There was a significant difference in HR between pelvic suspension group and control group on 15^th^ day. [C: 223.2±22.3/min vs HU: 279.9±0.75/min (p=0.031)]. (**Figure 4G)** Blood pressure responses to different doses of phenylephrine (5, 10, 15 & 20 µg/kg) and sodium nitroprusside (6, 12, 24 & 48 µg/kg) were recorded. There was significant lower blood pressure elevation to different doses of epinephrine in the pelvic suspension group as compared to control whereas significant lower blood pressure falls to different doses of nitroprusside in the pelvic suspension group. **(Figure 5)** We also found significant low BRS in pelvic suspension group as compared to control rats. BRS at 15^th^ day [C: 0.4957(0.4801-1.311) vs Ex: 0.2429(0.0499-0.4462) (p=0.026)]. **(Figure 4H)**

**Figure 5:**
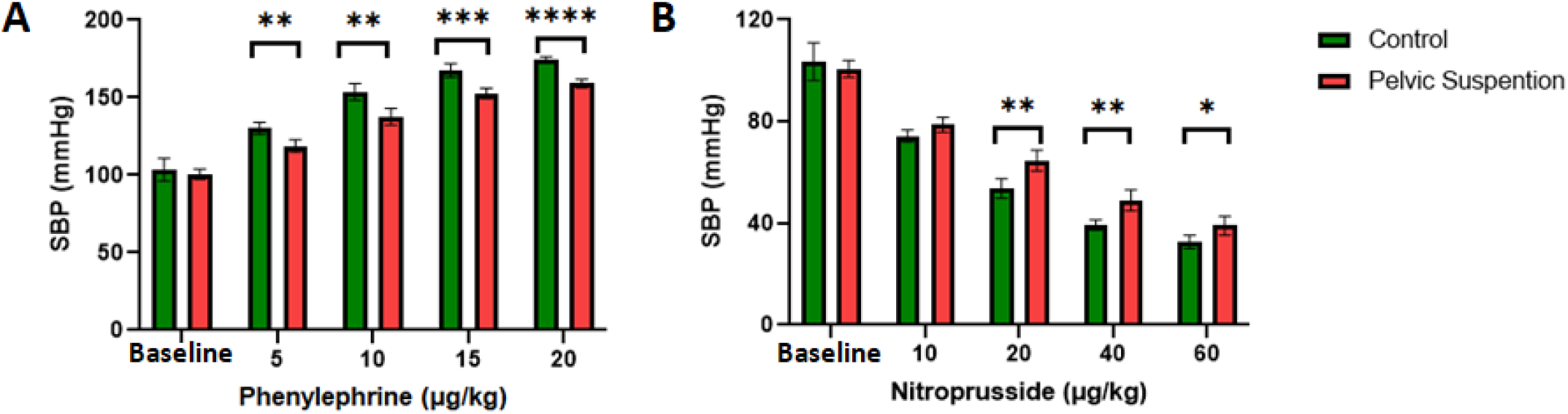
comparison of vasoconstriction and vasodilation property of vessels between control and pelvic suspension rats in vivo. **Figure5 A:** comparison of vasoconstrictive property of vessels (increase in mean arterial pressure) by injecting different dose of phenylephrine in carotid artery (5,10, 15 & 20 μg /kg) ; Two-way ANOVA showed significant effects of interaction F (4, 40) = 6.397 P=0.0004, Pelvic suspension F (1, 10) = 55.32 P<0.0001 and different dose of phenylephrine F (2.848, 28.48) = 600.9 P<0.0001. The Bonferroni post test showed significant differences between control and pelvic suspension at dose 0.1 μg/kg, 0.2 μg/kg, and 0.4 μg/kg; P< <0.05 * ; <0.01 **, <0.001 *** ; <0.0001 **** n=Control: 6-9 and pelvic suspension: 6-9, **Figure5 B:** comparison of Vasodilator property of vessels (fall in mean arterial pressure) by injecting different dose of nitroprusside in carotid artery (6, 12, 24 & 48 μg /kg); Two-way ANOVA showed significant effects of interaction F (4, 50) = 5.846 P=0.0006, Pelvic suspension F (1, 50) = 31.94 P<0.0001 and different dose of nitroprusside F (4, 50) = 553.7. The Bonferroni post test showed significant differences between control and pelvic suspension at dose 0.05 μg/kg, 0.1 μg /kg, 0.2 μg/kg and 0.5 μg/kg ; 0.0023**, 0.0020**, 0.0004***, <0.0001**** respectively. P< 0.05 *; <0.01 **, <0.001 ***, <0.0001 **** n=Control: 6-9 and pelvic suspension: 6-9,

To examine the structural changes in the carotid artery, H & E staining was done and the transverse section of the vessel was examined in different magnification **(Figure 2)**. Different comparative parameters such as Vessel wall thickness, luminal diameter, Intimal thickness & Smooth muscle layer thickness of both the groups of rats were assessed and measured. (Figure 4 I-L) show the Vessel wall thickness, luminal diameter, Intimal thickness & Smooth muscle layer thickness of both the groups of rats at the end of 15th day respectively. There was a trend of increase in the vessel wall thickness, Intimal thickness & Smooth muscle layer thickness in the pelvic suspension group but it was not significant; similarly non-significant reduction in luminal diameter in pelvic suspension group was observed. Vessel wall thickness [C: 125±23.34 µm vs HU: 146.5±12.05 µm (p=0.072)], luminal diameter [C: 2046±374.1µm vs HU: 1913±235.2 µm (p=0.475)], intimal thickness [C: 39.22±15.98 µm vs HU: 44.84±8.991 µm (p=0.47)], Smooth muscle layer thickness [C: 96.97±24.71µm vs HU: 114.5±17.88µm (p=0.189)] **(Figure 4 I-L)**

To examine the vasodilator and vasoconstriction properties of the carotid artery, cannulation was done and different doses of epinephrine and nitroprusside were used. Fig. 3 shows the change in the mean arterial blood pressure for the different doses of epinephrine and nitroprusside respectively. [Epinephrine 5 µgm/kg (C: 26.82±4.74 ±4.74 vs HU:18.21±6.21), 10 µgm/kg (C:50.26±7.01 vs HU:37.06±7.38), 15 µg/kg (C:63.97±5.9 vs Ex:52±4.3), 20 µg/kg (C:71.21±7.4 vs HU: 58.51±4.7) (p=0.0001)], Nitroprusside 10 µg/kg (C: -30.33±7.22 vs HU:-22 ±5.5), 20 µg/kg (C:50.26±5.25 vs HU:-36±6.16), 40 µg/kg (−65.7±7 vs -51.6±6.66), 60 µg/kg (E:-72±64 vs HU: -61.66±6.4) (p=0.0001)]. **(Figure 5A, B)**

Electron microscopy was also used to analyze structural alterations in the carotid artery microstructure, and the transverse section of the vessel was examined under different magnifications of 570X and 2250X. (Fig.5) Different comparison measures, such as endothelial layer, internal elastic lamina, smooth muscle, and external elastic lamina thickness, were assessed in both groups of rats. No significant change was observed in electron microscopy parameters between the control and pelvic suspension group. Endothelial layer thickness [C: 0.8556±0.4537µm vs HU: 0.8109±0.3848 µm (p=0.872)], internal elastic lamina [C: 2.044 ±0.6085 µm vs HU:1.685 ±0.5148 µm (p=0.1035), smooth muscle layer [C: 12.60 ±2.658 µm vs HU: 12.86 ±2.983 µm] and external lamina thickness [C: 4.327 ±0.6400 µm, Vs HU:4.354 ±1.019 µm (p=0.9152)]. **(Figure 3)**

Changes in the BRS were not accompanied by changes vascular structure, so it might be due to alteration in sympathetic and parasympathetic outflow. So we planned to compare autonomic tone of different group of rats by heart rate variability. To examine the effect chronic HU on sympathetic and parasympathetic system, we have done the HRV of both the group of rats. After 30 min of lead 2 ECG recording, we found significant change in R-R interval, SDSD, RMSD, VLF and HF power and LF/HF ratio between control and HU group after 15 days of HU.. Different parameters were compared such as R-R interval, SDSD, SDRR, RMSD, Total power, VLF, LF< HF and LF/HF ratio. We found significant change in R-R interval, SDSD, RMSD, VLF and HF power and LF/HF ratio between control and HU group after 15^th^ days of HU. R-R interval [C: 158.33±7.17 vs HU:149.67±3.61(p=0.0246)], SDSD [C: 7.73±0.18 vs HU: 3.6±0.29 (p=0.0001), SDRR [C: 7.1±1.7 vs HU:7.6±1.5 (p=0.6064), RMSD [C: 7.63±0.43 vs HU: 3.46±0.31 (p= 0.0001), Total power [C: 47.30±17.63 vs HU: 60.31±4.31 (p= 0.119), VLF [C: 38.99±4.22 vs HU: 9.05±3.06 (p= 0.001), LF [C: 10.64±1.42 vs HU: 24.44±9.76 (p= 0.0054), HF [C: 24.44+9.76 vs HU: 10.23±1.38 (p= 0.0054), LF/HF Ratio [C: 0.39±0.13 vs HU: 1.05±0.18 (p= 0.0001).**(Figure 6)**

**Figure 6:**
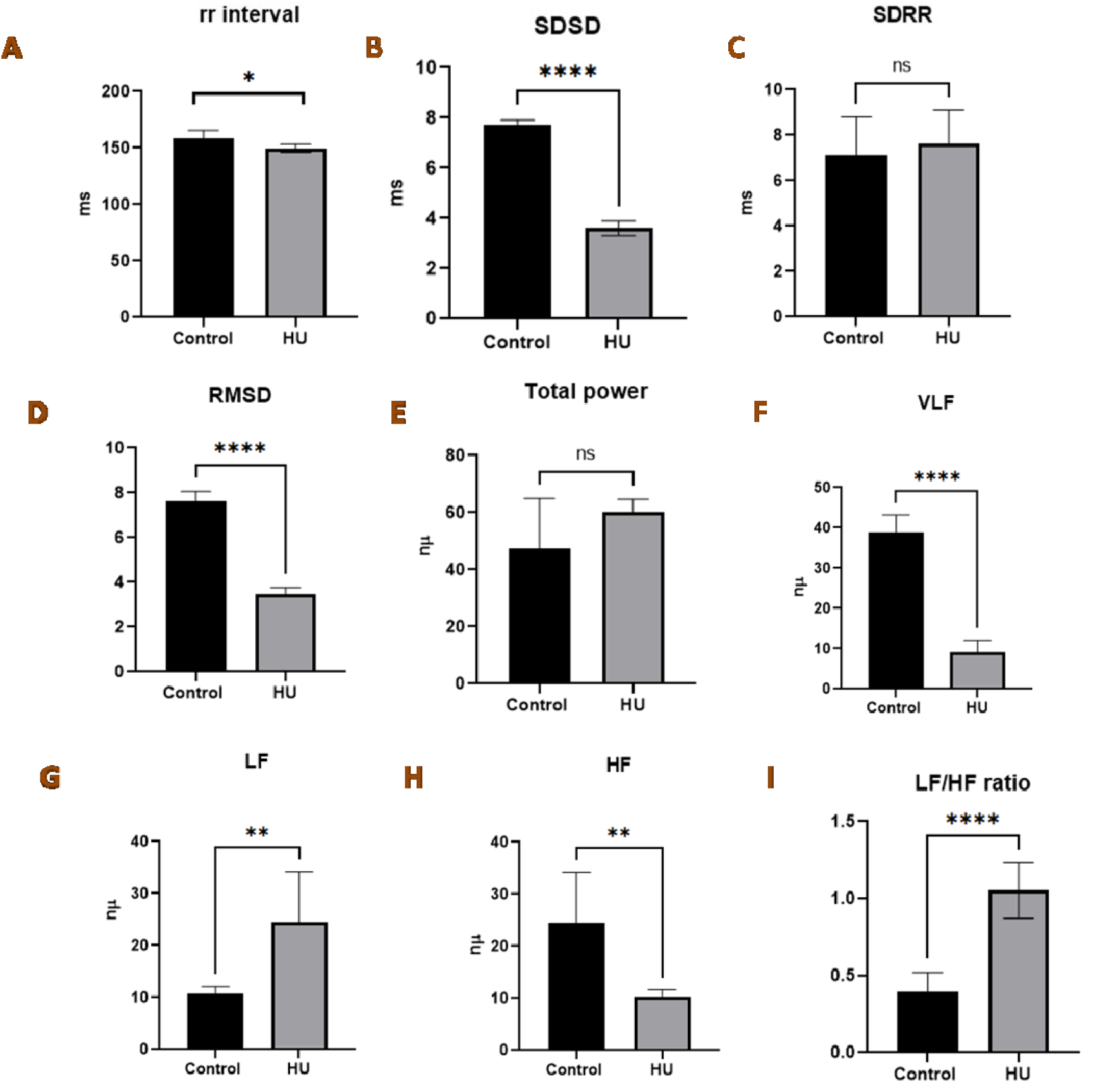
comparison of Heart rate variability parameters such as rr interval, SDSD, SDRR, RMSD, Total power, VLF, LF, HF, LF/HF ratio between control and pelvic suspension rats after 15 days. **Figure6 A:** *comparison of* rr interval (ms) of pelvic suspension rats with control rats after 15 days of HU. Unpaired T test; 2 tailed 95% confidence interval, [C: 158.33±7.17 vs HU:149.67±3.61(p=0.0246)] n=control:6-9 and pelvic suspension:6-9 **Figure 6 B :** *comparison of* Heart rate variability parameters ; SDD of pelvic suspension rats with control rats after 15 days of HU. Unpaired T test; 2 tailed 95% confidence interval, [C: 7.73±0.18 vs HU: 3.6±0.29 (p=0.0001) n=control:6-9 and pelvic suspension:6-9 **Figure 6 C :** *comparison of* Heart rate variability parameters ; SDRR of pelvic suspension rats with control rats after 15 days of HU. Unpaired T test; 2 tailed 95% confidence interval, [C: 7.1±1.7 vs HU:7.6±1.5 (p=0.6064), n=control:6-9 and pelvic suspension:6-9 **Figure 6D:** *comparison of* Heart rate variability parameters; RMSD pelvic suspension rats with control rats after 15 days of HU. Unpaired T test; 2 tailed 95% confidence interval, [C: 7.63±0.43 vs HU: 3.46±0.31 (p= 0.0001), n=control:6-9 & pelvic suspension:6-9 **Figure 6E:** *comparison of* Heart rate variability parameters; total power of pelvic suspension rats with control rats after 15 days of HU. Unpaired T test; 2 tailed 95% confidence interval, [C: 47.30±17.63 vs HU: 60.31±4.31 (p= 0.119)] n=control:6-9 and pelvic suspension:6-9 **Figure 6F:** *comparison of* Heart rate variability parameters; VLF of pelvic suspension rats with control rats after 15 days of HU. Unpaired T test; 2 tailed 95% confidence interval, [C: 38.99±4.22 vs HU: 9.05±3.06 (p= 0.001), n=control:6-9 & pelvic suspension:6-9 **Figure 6G:** *comparison of* Heart rate variability parameters; LH of pelvic suspension rats with control rats after 15 days of HU. Unpaired T test; 2 tailed 95% confidence interval, [C: 10.64±1.42 vs HU: 24.44±9.76 (p= 0.0054), n=control:6-9 and pelvic suspension:6-9 **Figure 6H:** *comparison of* Heart rate variability parameters; HF pelvic suspension rats with control rats after 15 days of HU. Unpaired T test; 2 tailed 95% confidence interval, [C: 24.44+9.76 vs HU: 10.23±1.38 (p= 0.0054), n=control:6-9 and pelvic suspension:6-9 **Figure 6I:** *comparison of* Heart rate variability parameters; LF/HF ratio of pelvic suspension rats with control rats after 15 days of HU. Unpaired T test; 2 tailed 95% confidence interval, 0.39±0.13 vs HU: 1.05±0.18 (p= 0.0001).,n=control:6-9 and pelvic suspension:6-9 P< 0.05 *; <0.01 **, <0.001 ***, <0.0001 **** n=Control: 6-9 and pelvic suspension: 6-9 Heart rate variability (HRV), Standard deviation of successive RR interval differences (SDSD), standard deviation of the RR interval (SDRR), Root mean square of successive differences (RMSSD), Very low frequency (VLF), Low Frequency, High frequency (HF)

## Discussion

We have simulated chronic microgravity in a rat model using the pelvic suspension method. We found significant changes in body weight, food intake, water intake, HR, BRS, vasodilator, and vasoconstrictor responses.

The body weights of the HU rats were significantly low as compared to the control rats. We found that the reduction in body weight is a direct consequence of the decrease in food intake. A similar drop in body weight was documented in earlier experiments of tail suspension rats. (12) A recent study by Jin et al. (2018) showed that simulated weightlessness can modify intestinal permeability, which could also contribute to reduction in body weight (15). As the suspension period came to an end, food and water intake showed a trend toward increasing. This could be attributed to the habituation of rats after a few days of suspension with the pelvis brace. Our experiments demonstrated significant changes in water intake in pelvic-suspended rats after 15 days of suspension. Many studies have compared the water intake after HU, however the results are inconsistent. Steffen et al. (1990) showed a significant decrease in water intake, while Martel et al. (1996) found no significant reduction in water intake after HU. (16, 17)

We found no significant difference in hemodynamic parameters such as SBP, DBP, and MAP in the HU group against the control group. Human (18) and rat (19) investigations revealed similar results. In our study, we found that the heart rate of the pelvic suspended group increased significantly. Human investigations conducted during Spacelab missions revealed that after 15 days of space flight, heart rates were raised due to microgravity exposure on the day of landing and three days after landing (20). Analogously, Moffitt et al. showed that a 14-day hindlimb suspension increased heart rate not just at 14 day time but also at the early and midpoints (21).

We found a significant decrease in the BRS in the pelvic suspension group as compared to the control group. Although this outcome is consistent with some of the previous research (5-6), no effect has also been reported in other experiments (22-26). This might be the result of a suspension with a low degree or fewer days. A smaller range of baroreflex buffering suggests a possible decrease in the capacity to increase heart rate via altering sympathetic reactivity or vagal tone. Reductions in the responsiveness of the carotid baroreceptor-reflex have been seen during 4-to 10-day space shuttle flight (27). Ambulatory blood pressure recordings revealed that BRS was decreased in astronauts both during and following the flight, (28) whereas other reports indicate that there was no postural hypotension because there was no change in the vasodilator and vasoconstrictor responses to postural variation during the mission. (29)

We found an increasing trend in the vessel wall thickness, Intimal thickness, smooth muscle layer thickness and the luminal diameter but these changes were non-significant changes. Gao et al. (2009), revealed similar results such as increase in the media thickness of the common carotid artery and basilar artery after 28 days of tail suspension. (30) In some of the previous studies, vascular adaptation to microgravity has been suggested to be a generalized or universal response, whereas others suggest regional phenomena. Pressure may be the predominant trigger causing artery regional adaptation. The redistribution of transmural pressure across the vasculature appears to be the primary stimulus for vascular adaptation to microgravity, with flow and volume redistribution following as a secondary consequence. Even when the body fluid redistribution has found a new equilibrium, the pressure redistribution remains as long as the head-down tilt or microgravity exposure is maintained. (31)

The *comparison of* HRV between HU and control rats revealed a significant difference in sympathovagal tone, which could be the cause of tachycardia in HU rats at rest. However, this change in sympathovagal balance had no significant effect on mean arterial pressure as many previous studies had suggested. Moffitt et al. confirmed impaired sympathovagal balance in HU rats by reporting a larger decrease in heart rate in HU rats by Propranolol-induced cardiac sympathetic inhibition and no significant rise in heart rate above baseline in HU rats by methylatropine-induced parasympathetic inhibition. They also found that the heart rate response to propranolol was much higher after atropine therapy in the CC group.(32) Collectively results of our study suggest 15 days of hind limb suspension leads to increase sympathetic tone and decrease in parasympathetic tone. We also found impairment in VLF power in HRV which suggests impairment in rennin angiotensin Aldosterone system in HU. (33)

Many investigations have found that microgravity exposure leads to decreased parasympathetic and increased sympathetic cardiac tone in astronauts. The rise in sympathetic tone in this study may have been caused solely by chronic stress, but this is not the case as HU does not produce stress, as reported by Moffitt et al., after 15 days of HU, the levels of cortisol in hind limb unloading and control rats were same. (32) We found no significant association between BRS and vascular structural parameters, implying that changes in BRS occur prior to changes in carotid artery structure by pelvic suspension.

Based on these data, we propose that the persistent cephalic shift of body fluid in the blood vessels above the heart leads to resetting of the baroreceptor operation point and range, which leads to an impaired baroreceptor response in HU rats. Impaired baroreceptor response is due to impaired sympathovagal after 15 days of pelvic suspension. If the exposure to microgravity continues for a longer period of time, the above phenomena become more stable, and the muscle cells in the carotid arteries, which are constantly subjected to a higher pressure, may show hypertrophy to compensate for the high pressure that we were unable to obtain due to short duration of intervention.

## Conclusion

We observed that 15 days of microgravity simulation using pelvic suspension reduced baroreceptor sensitivity, but not enough to cause anatomical changes in the carotid artery. Changes in the BRS are accompanied by changes in the vascular vasoconstriction and vasodilation properties of HU rats, attributable to changes in sympathetic and parasympathetic output.

